# Natural *Wolbachia* infections are common in the major malaria vectors in Central Africa

**DOI:** 10.1101/343715

**Authors:** Diego Ayala, Ousman Akone-Ella, Nil Rahola, Pierre Kengne, Marc F. Ngangue, Fabrice Mezeme, Boris K. Makanga, Carlo Costantini, Frédéric Simard, Franck Prugnolle, Benjamin Roche, Olivier Duron, Christophe Paupy

**Author notes:** Corresponding author: Diego Ayala, MIVEGEC, IRD, CNRS, Univ. Montpellier, 911 av Agropolis, BP 64501, 34394 Montpellier, France.

## Abstract

During the last decade, the endosymbiont bacterium *Wolbachia* has emerged as a biological tool for vector disease control. However, for long time, it was believed that *Wolbachia* was absent in natural populations of *Anopheles.* The recent discovery that species within the *Anopheles gambiae* complex hosts *Wolbachia* in natural conditions has opened new opportunities for malaria control research in Africa. Here, we investigated the prevalence and diversity of *Wolbachia* infection in 25 African *Anopheles* species in Gabon (Central Africa). Our results revealed the presence of *Wolbachia* in 16 of these species, including the major malaria vectors in this area. The infection prevalence varied greatly among species, confirming that sample size is a key factor to detect the infection. Moreover, our sequencing and phylogenetic analyses showed the important diversity of *Wolbachia* strains that infect *Anopheles.* Co-evolutionary analysis unveiled patterns of *Wolbachia* transmission within *Anopheles* species, suggesting that past independent acquisition events were followed by co-cladogenesis. The large diversity of *Wolbachia* strains that infect natural populations of *Anopheles* offers a promising opportunity to select suitable phenotypes for suppressing *Plasmodium* transmission and/or manipulating *Anopheles* reproduction, which in turn could be used to reduce the malaria burden in Africa.

## Introduction

Malaria still affects millions of people and is the cause of thousands of deaths worldwide, although sub-Saharan Africa pays the highest tribute^1^. Currently, vector-control measures (e.g., insecticide-treated bed nets or indoor residual spray) are the largest contributors to malaria eradication^2^. If these interventions are maintained or increased, malaria burden should be drastically reduced in Africa before 2030^3^. These predictions are based on the constant effectiveness of these methods. However, the spread of insecticide resistance^4^ and vector behavioural changes related to the massive use of bed nets^5^ might challenge malaria eradication in the coming decades. Therefore, it is vital to develop alternative and environmentally friendly control strategies for the millennium development goal of malaria eradication^6^.

Several methods have been proposed to accompany or replace the use of synthetic insecticides^7^ Among them, the use of the maternally inherited *Wolbachia* bacteria (α-proteobacteria, Anaplasmataceae family) has emerged as a promising alternative biological tool for fighting malaria and other vector-borne diseases^7–11^. This bacterium exhibits a large spectrum of interactions with its hosts: from mutualism and commensalism to parasitism^12^ Moreover, *Wolbachia* can invade mosquito populations and/or prevent vector-borne infections in some of the most important mosquito vectors^8,11,13^. Indeed, *Aedes aegypti* populations that were artificially infected with *Wolbachia* have been successfully used to suppress dengue transmission in laboratory conditions, and have been released in natural populations of this mosquito^14,15^. Similarly, laboratory studies showed that infection of *Anopheles* (the vector of human malaria) with *Wolbachia* strains has a negative impact on the transmission of *Plasmodium* parasites^16–18^, providing a relevant alternative for malaria control. Unfortunately, only one stable transfected *Wolbachia* colony has been described in *Anopheles stephensi*^18^. Therefore, data on the use *Wolbachia* for *Anopheles* control remain scarce and mainly concern experimental studies in laboratory conditions^18,19^, due to technical (i.e., egg microinjection) and biological (i.e., competitive exclusion with the bacterium *Asaia)* difficulties to carry out transinfections in *Anopheles*, despite multiple assays^20–22^ For long time, it was believed that *Wolbachia* was absent in natural populations of *Anopheles*^20^. However, in the last few years, three studies reported that *An. gambiae, An. coluzzii* and *An. arabiensis* (three major malaria vectors) populations from Burkina Faso and Mali (West Africa) are naturally infected by *Wolbachia*^23–25^. Noteworthy, they showed a negative correlation between *Wolbachia* infection and *Plasmodium* development^24,25^. Moreover, a very recent report suggests that other *Anopheles* species also are infected with *Wolbachia^22^.* These findings support the development of novel vector control strategies based on *Wolbachia-Anopheles* interactions. However, although *Wolbachia* naturally infects 40% – 60% of arthropods^26,27^, infection of *Anopheles* species is still not well documented. Moreover, during the last decade, screens in many other malaria mosquito species worldwide (n=38) did not bring any evidence of *Wolbachia* infection^9,20,28^.

In this study, we investigated the presence of *Wolbachia* in 25 *Anopheles* species in Gabon, Central Africa. We sampled mosquitoes across the country and in a variety of ecological settings, from deep rainforest to urban habitats. By using a molecular approach, we confirmed *Wolbachia* presence in 16 species, including all the major malaria vectors in Central Africa *(An. gambiae, An. coluzzii, An. funestus, Anopheles nili* and *Anopheles moucheti*). The prevalence of *Wolbachia* infection was particularly high in *An. nili* and *An. moucheti.* Phylogenetic analysis revealed that all the infected mosquito species hosted *Wolbachia* bacteria belonging to the supergroup A or B (both exhibit high genetic diversity). Finally, we explored the co-evolution between *Wolbachia* and *Anopheles.* The results have direct implications for the development of new and environmentally friendly vector control strategies and open new directions for research on pathogen transmission and reproductive manipulation.

## Results

### Wolbachia naturally infects a large number of Anopheles species from Gabon

In this study, we screened 648 mosquitoes from eight sites in Gabon (Fig. 1, Table 1, Table S1). On the basis of their morphological traits^29^ and molecular analysis results^30–34^, we identified 25 *Anopheles* species (Text S1). Our sample included all the species in which the presence of *Wolbachia* was previously investigated in Africa (An. *gambiae, An. coluzzii, An. funestus* and *An. coustani*), with the exception of *An. arabiensis* that is absent in Gabon (Table 1)^35^. By PCR amplification of a *16S* rRNA fragment^24^, we found 70 *Wolbachia*-positive specimens that belonged to 16 different *Anopheles* species, distributed throughout the country (Fig. 1, Table S1). With few exceptions, *Wolbachia* infection rate was lower than 15% in most species (n=11), as observed in other arthropods^26,27^ On the other hand, for *An. moucheti, An. “GAB-2”, An. “GAB-3”, An. jebudensis* and *An. nili*, more than 50% of sampled mosquitoes were infected (Table 1), as previously reported in other mosquito species where prevalence can be very high^36,37^ To our knowledge, none of these species was previously screened for *Wolbachia* infection.

**Figure 1.**
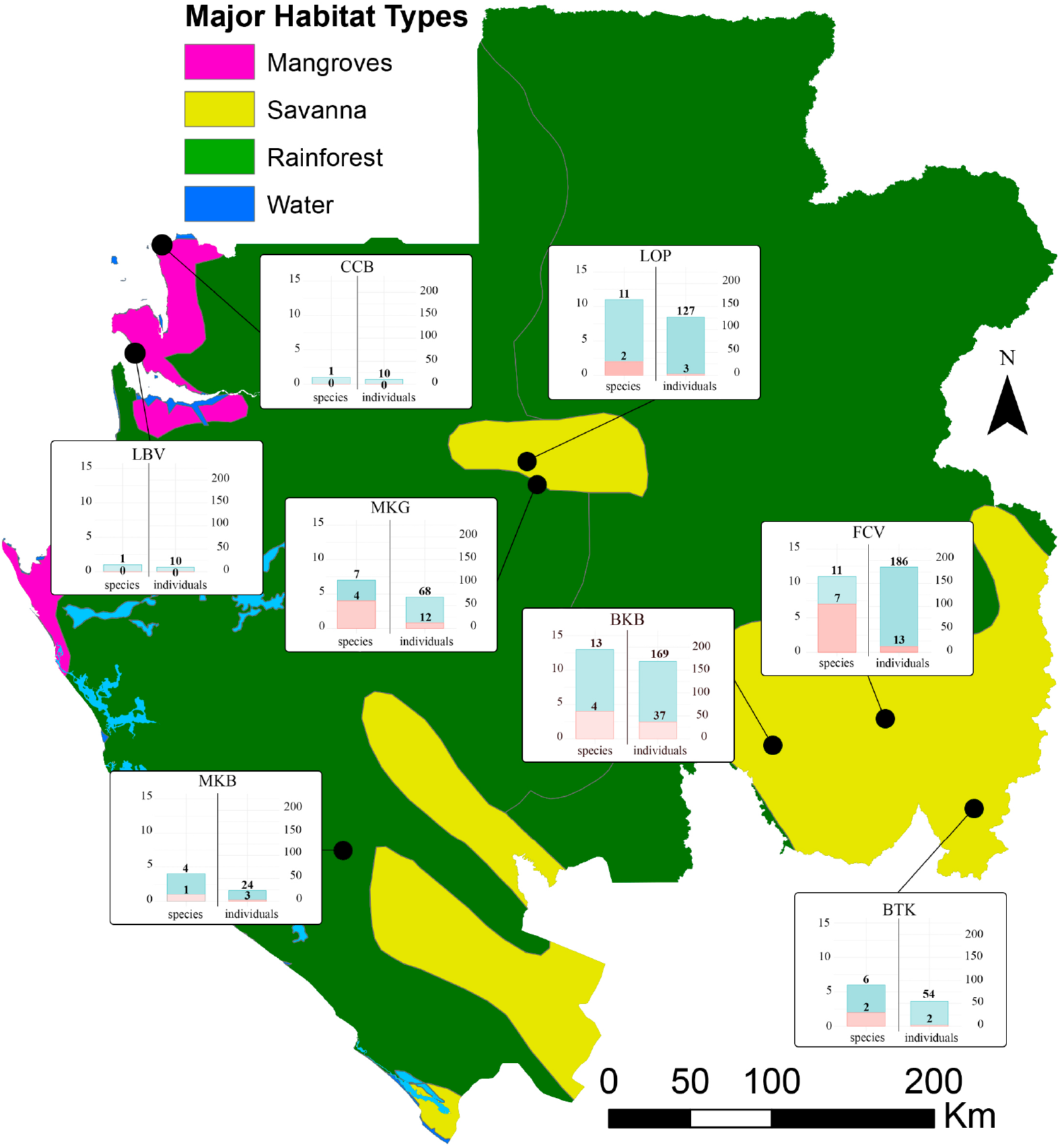
Sampling sites and *Wolbachia* infection prevalence. Map of Gabon showing the main African habitat types (^78^, freely available at http://maps.tnc.org/gis_data.html) and the villages were sampling took place (black dots). The map was drawn using ArcGIS Basic v.10. The prevalence of *Wolbachia* infection (number of infected *Anopheles* species and individuals) per site is presented in bar charts. The pink colour indicates positive species/individuals and blue the total number of species/individuals screened for *Wolbachia* infection at that site. CCB: Cocobeach; LOP: Lopé; MKG: Mikongo; BTK: National Park of Plateaux Batékés; FCV: Franceville; LBV: Libreville; MKB: National Park of Moukalaba-Doudou; BKB: Bakoumba.

**Table 1.**
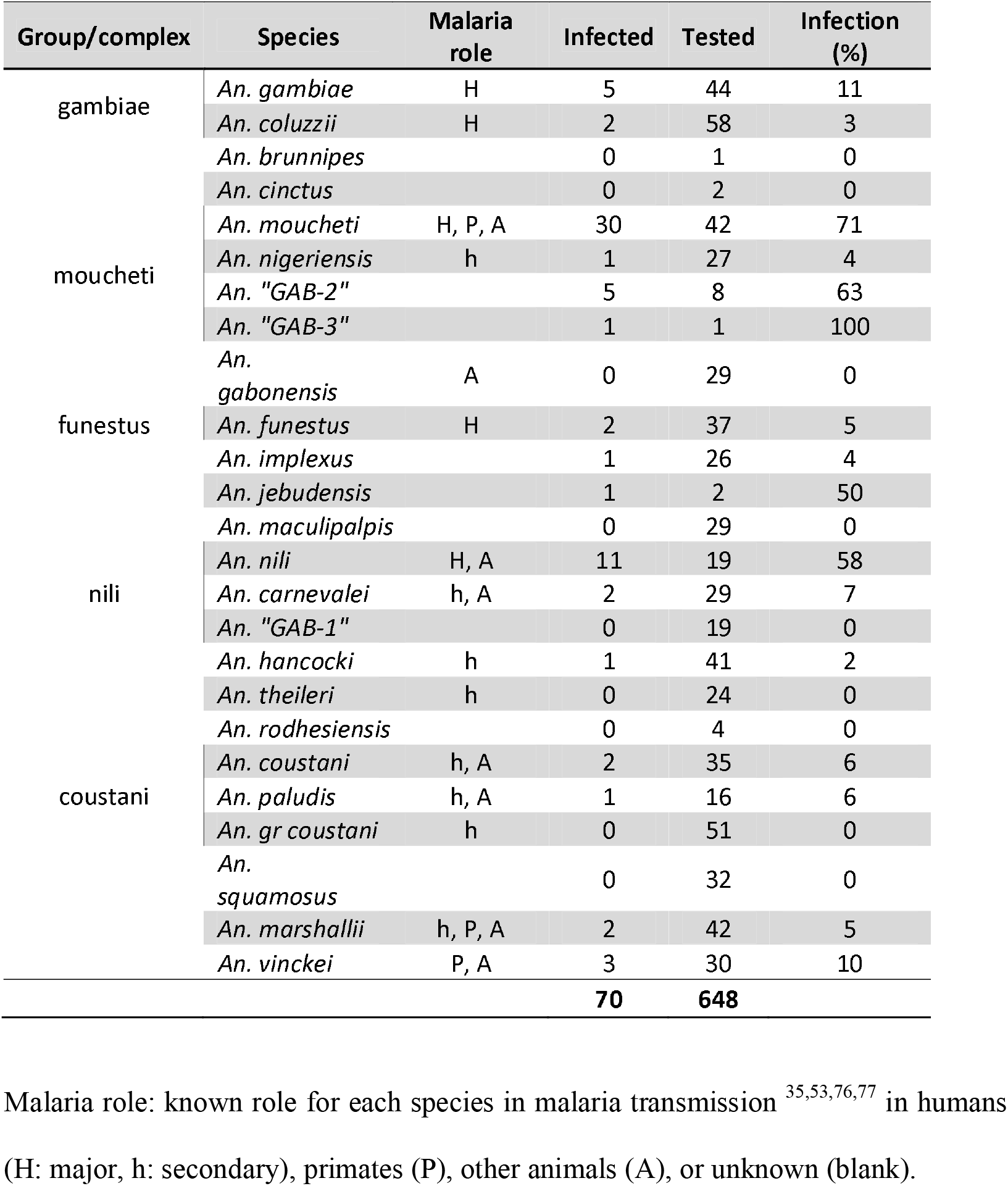
Summary of the *Anopheles* species screened in this study.

### Wolbachia is maternally inherited in An. moucheti

Although *Wolbachia* is mainly maternally transmitted^12^, horizontal transmission may occasionally occur in natural conditions^38–40^. To confirm the maternal transmission in the infected mosquito species, we focused on *An. moucheti* for logistic reasons (i.e., highest *Wolbachia* prevalence and ease of sampling). Although no laboratory *An. moucheti* strain is currently available, we obtained eggs from six *Wolbachia-infected* females. In total, we analysed the infectious status of 79 progeny by PCR amplification of the same *16S* rRNA fragment^24^ (Table S2), and found that 70 were infected, with an average maternal transmission frequency of 97.54% (range: 90% – 100%).

### Naturally occurring *Wolbachia* strains in *Anopheles* reveal high genetic diversity

By sequence analysis of the *16S* rRNA fragment PCR-amplified from each *Anopheles* sample (Table 1), we could assign the *Wolbachia* strains to three pre-existing supergroups: A (n=5), B (n=64) and C (n=1) (Fig. 2). Specifically, we detected supergroup B *Wolbachia* in 64 mosquitoes belonging to all 16 infected *Anopheles* species. We found supergroup A *Wolbachia* in five individuals from four species (*An. funestus, An. coluzzii, An. vinckei* and *An. carnevalei)*, thus providing examples of multiple infections, as previously observed in *Ae. albopictus*^41^ (Fig. 2). None of the mosquitoes examined was co-infected by *Wolbachia* strains belonging, for instance, to the supergroups A and B. Moreover, we confirmed that the *Wolbachia* strains previously identified in *An. gambiae s.l.* from Burkina Faso and Mali are included in the supergroups A and B^23,25^. Finally, we found that one *An. coustani* individual was infected by a *Wolbachia* strain from supergroup C that is known to infect only filarial worms. Therefore, we investigated the presence of filarial nematode DNA in the mosquito by PCR amplification and sequencing of a fragment of the *COI* filarial gene^42^, followed by phylogenetic analysis with RAxML. Our results confirmed the presence of *Dirofílaria immitis* in this specimen (Fig. S1). This canine filarial parasite hosts *Wolbachia* and is transmitted by many mosquitoes, including *Anopheles*^43^. Therefore, it is not surprising to find an *An. coustani* specimen infected by this filarial nematode. This specimen was excluded from further investigations.

**Figure 2.**
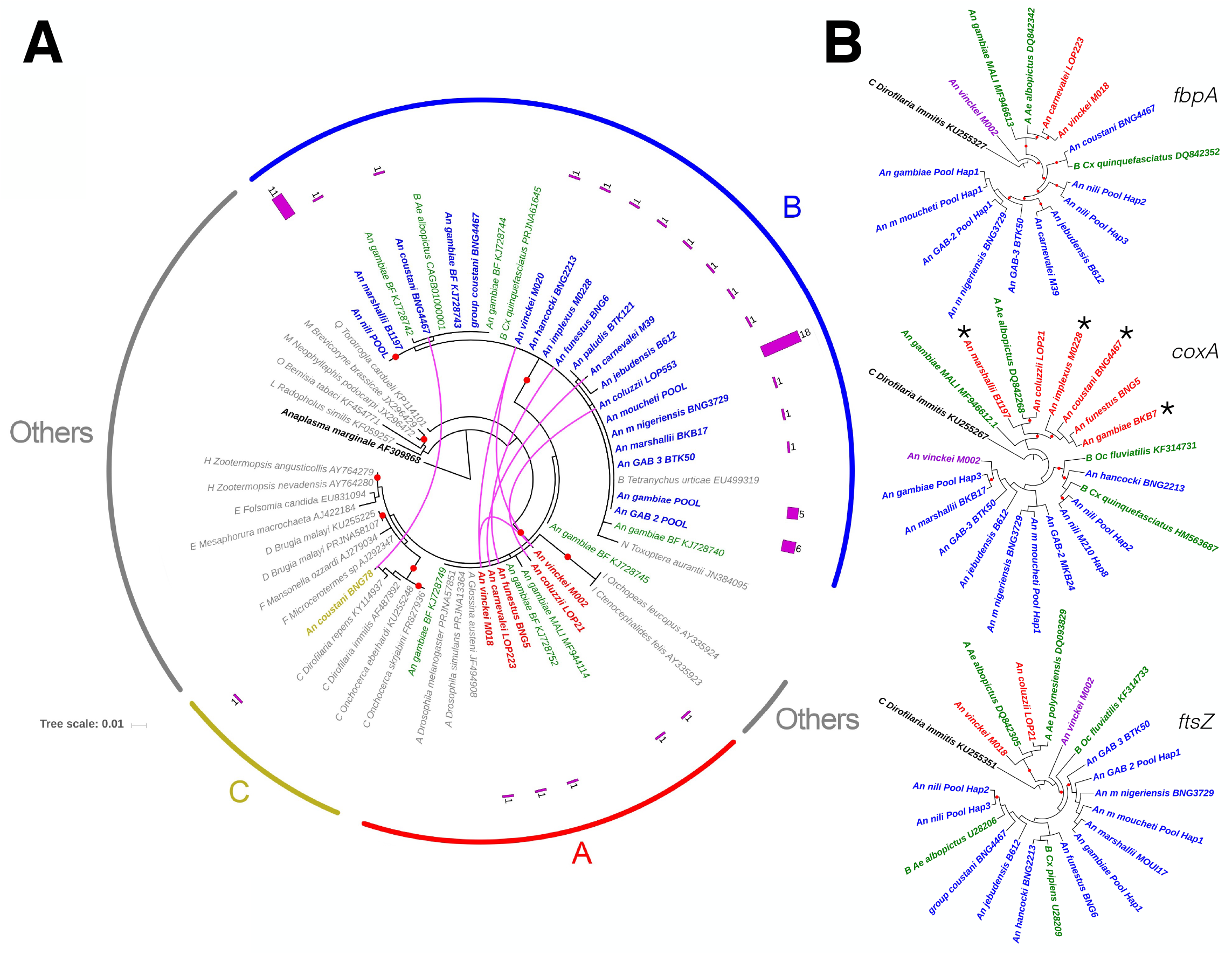
Circular phylograms of the *Wolbachia* strains isolated in the 16 *Anopheles* species. The phylogenetic trees were built with RAxML^72^ The names of the *Anopheles* species from which the *Wolbachia*-specific sequences were isolated in this study are shown in blue (positive for *Wolbachia* supergroup B), red (positive for supergroup A) and brown (positive for supergroup C), while the names of mosquitoes species (*Diptera*) from which the previously published *Wolbachia* sequences were isolated are in green. Red dots show branches supporting a bootstrap >70% from 1000 replicates. (A) Circular phylogenetic tree using the *Wolbachia*-specific *16S* rRNA fragment and *Anaplasma marginale* as outgroup. Different *Wolbachia* strains found in the same *Anopheles* species are connected by pink lines. The pink bar charts indicate the number of identical *Wolbachia* haplotypes found in each species. Scale bar corresponds to nucleotide substitutions per site. (B) Circular phylogenetic trees based on the *coxA, fbpA and ftsZ* fragment sequences using *Dirofilaria immitis* (supergroup C) as outgroup. Specimens with a different supergroup assignation than *16S* are marked with asterisks.

To expand our knowledge on the *Wolbachia* strains that infect natural *Anopheles* populations, we PCR amplified, sequenced and analysed a *16S* rRNA fragment and also fragments from three other conserved *Wolbachia* genes *(coxA, fbpA* and *ftsZ)* that are commonly used for strain typing and evolutionary studies^44^ (Fig. 2). We used a new nested PCR protocol (see *Methods)* for samples that could not be genotyped using the classical Multi-Locus Sequence Typing (MLST) primers (Table S1). Our phylogenetic analyses confirmed the *16S* results, assigning most of the species to supergroups A and B. Few samples (asterisks in Fig. 2, gene *coxA*) showed some incongruence relative to the *16S* results. They suggest signals of recent recombination between the supergroups A and B, as previously demonstrated^44^ Detailed sequence analysis revealed that mosquito species belonging to the same group or complex (i.e., *An. moucheti* and *An. gambiae)* displayed a common *Wolbachia* haplotype (defined here as a unique allelic profile) (Fig. 2 and Fig. 3). Conversely, some species with lower prevalence (i.e., *An. coluzzii, An. marshallii, An. vinckei* or *An. funestus)* displayed a variety of haplotypes. The case of *An. vinckei* was particularly interesting because the three infected specimens displayed different haplotypes for the analysed *Wolbachia* genes. Moreover, one specimen (An. *vinckei* M002, Fig. 2) was infected by a completely different *Wolbachia* strain. Overall, the *Wolbachia* haplotypes identified in this study were different from the allelic profiles of the previously annotated *Wolbachia* strains or of the strain that infects *An. gambiae* in Burkina Faso and Mali^23,25^ (Fig. 2 and Fig. 3). Within supergroup B, we could easily distinguish at least two strains. The strain infecting *An. moucheti (wANMO)* was similar to the one identified in *An. gambiae* (in our study) or *An. marshallii*, while the strain infecting *An. nili (wANNI)* was more closely related to those found in other mosquito species, such as *Ae. albopictus* or *Cx. quinquefasciatus* (Fig. 2 and Fig. 3). Conversely, the other haplotypes were associated with one specific host.

**Figure 3.**
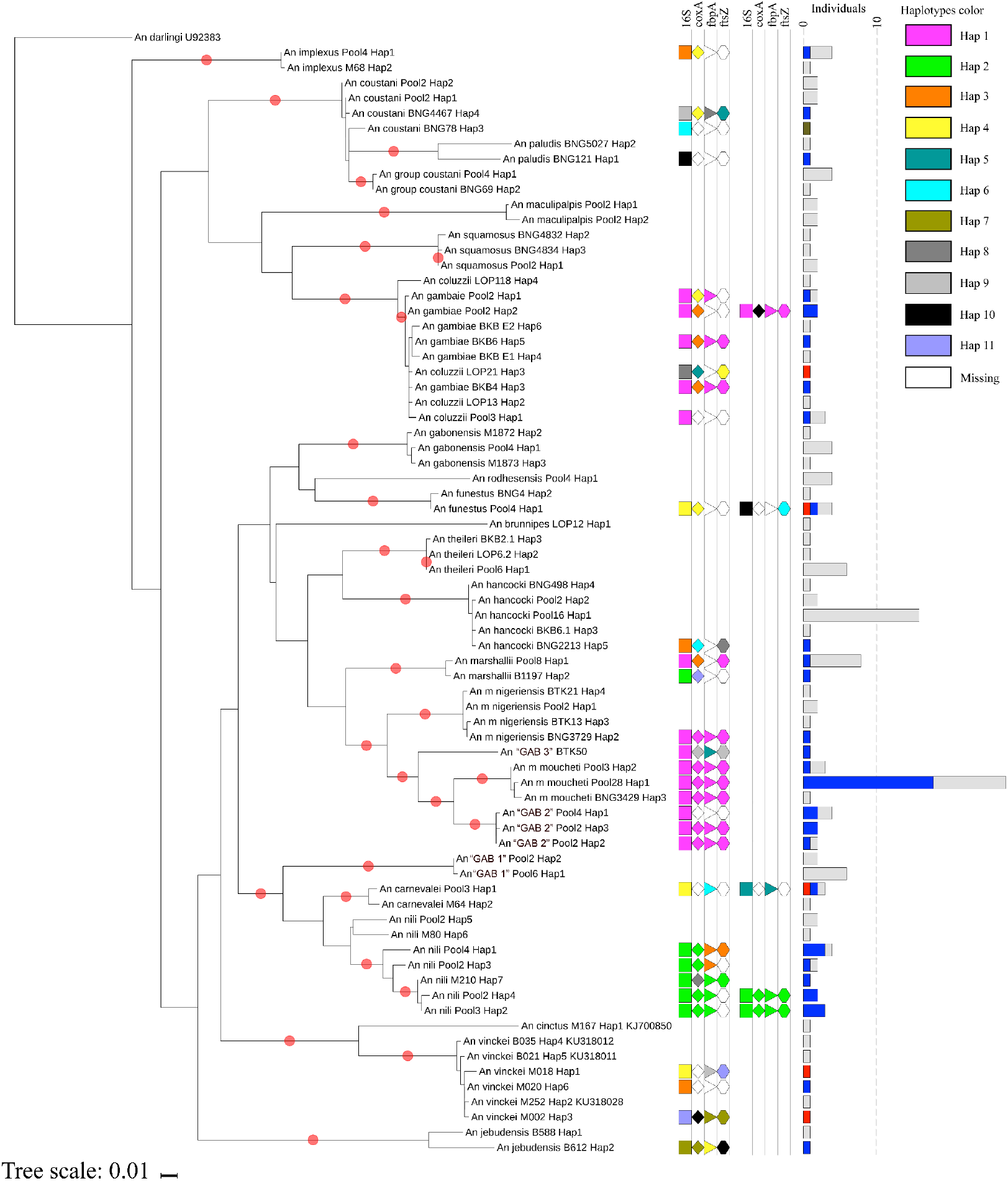
Maximum likelihood phylogeny of the 25 *Anopheles* species under study and *Wolbachia* haplotypes. The tree was inferred with RAxML^72^ using the sequences of the *COII* fragment from 176 *Anopheles* specimens belonging to the 25 species under study and rooted with *Anopheles darlingi* as outgroup (New World mosquito, diverged 100 Myr ago^79^). Red dots in branches represent bootstrap values >70% from 1000 replicates. The shape of each field column represents the *16S* (rectangle), *coxA* (rhombus), *fbpA* (triangle) and *ftsZ* (hexagon) genes. The different *Wolbachia* gene haplotypes (i.e., unique allelic profiles) are indicated with colour codes (all pink = the newly identified wANMO strain). The bar chart size indicates the number of individuals of the same species with the same haplotype, and the colour represents their infection status: grey, noninfected; blue, infected by the *Wolbachia* supergroup B; red, infected by supergroup A; brown, infected by supergroup C.

### *Wolbachia* independently evolves in malaria-transmitting mosquitoes

As *Wolbachia* is mainly a maternally inherited bacterium, the host mitochondrial DNA (mtDNA) is a suitable marker to study its evolutionary history in *Anopheles*^45^. Analysis of *COII* sequences from 176 specimens belonging to the 25 *Anopheles* species collected in Gabon provided the most exhaustive phylogenetic tree of *Anopheles* in Central Africa (Fig. 3). This analysis highlighted the independent acquisition and loss of *Wolbachia* across the different *Anopheles* species clades. Moreover, the genetic distances of *Wolbachia* strains and their *Anopheles* host were not correlated (Mantel test, p >0.05) (Fig. S2). Nevertheless, mosquitoes from the same complex, and therefore genetically very close, shared the same *Wolbachia* supergroup and haplotypes (Fig. 3 and Fig. S2). Finally, we investigated how *Wolbachia* evolved within each *Anopheles* species^46^. Our results revealed that *Wolbachia*-infected and non-infected mosquitoes shared the same mtDNA haplotype (Fig. 2), indicating that infection status and host haplotypes are not associated.

## Discussion

The present study provides three key conclusions. First, the genus *Anopheles* includes a large number of species that are naturally infected by *Wolbachia* (16/25), with high infection prevalence among major malaria vectors. Second, *Anopheles-infecting Wolbachia* bacteria show high genetic diversity, with similar haplotypes detected in different *Anopheles* species. Third, the independent evolution of *Wolbachia* and *Anopheles* might be interpreted as multiple acquisition events with horizontal transmission. The large diversity of *Wolbachia* strains that infect many natural *Anopheles* populations could represent a major opportunity for reducing pathogen transmission and/or for reproductive manipulation in *Anopheles* with the aim of decreasing malaria burden in Africa.

For long time, the scientific community has been interested in finding new ways to use *Wolbachia* for fighting vector-borne diseases^7–9,11^. In arthropods, *Wolbachia* is broadly spread, including among *Culex* and *Aedes* mosquitoes. Conversely, it was believed that in the genus *Anopheles, Wolbachia* infections were mainly limited to species within the gambiae complex^23,25^, and few other species^22^. Several hypotheses can be put forward to explain this. First, low infection prevalence or local variations could have hindered the discovery of *Wolbachia* infections. In our study, most *Anopheles* species exhibited a prevalence lower than 15% (Table 1). This pattern is common in many other arthropods^26,27^, and it is usually associated with a weak manipulation of the host reproduction and/or imperfect maternal transmission^47^ However, high *Wolbachia* prevalence rates have been reported for some mosquitoes species, such as *Culex pipiens*^36^ or *Aedes albopictus^48^*. In general, our sampling effort was higher than in previous studies (n <30)^9,28^, and this could explain why we found more infected species. Our statistical analysis showed that a sample size of 60 individuals per species is needed to quantify correctly prevalence rates lower than 15%, with a probability of 95% (Fig. S3). Moreover, local frequency variations among populations could also hinder the detection of *Wolbachia* infections^36^. For instance, we sampled *An. coluzzii* in three different sites, but we only found *Wolbachia-infected* mosquitoes at La Lopé (Fig 1, Table S1). Therefore, sampling in different localities and in different seasons might improve detection rates. Second, it could be difficult to detect low-density *Wolbachia* infections in *Anopheles* with the routinely used molecular tools, as previously reported for other arthropods^49,50^ and recently in *Anopheles gambiae*^25^. Our results indicate that conventional PCR amplification (wsp-targeting primers^44^) analysis allowed detecting *Wolbachia* infection only in 6 of the 16 species *(An. moucheti, An m. nigeriensis, An. “GAB-3”, An nil, An. jebudensis* and *An. vinckei)* under study, presumably because of the high *Wolbachia* density. Moreover, some *Anopheles* species with high *Wolbachia* infection rates, such as *An. moucheti* or *An. nili*, were never screened before.

Unexpectedly, our work revealed that *Anopheles* species are infected by different *Wolbachia* strains. Although three previous studies reported *Wolbachia* infection in the *An. gambiae* germline^23–25^, we do not know whether *Wolbachia* naturally invades and is maternally inherited in the infected mosquito species from Gabon. Indeed, horizontal gene transfer (resulting in the insertion of *Wolbachia* genes within the mosquito genome), or parasitism (e.g., by filarial nematodes) could explain the detection of *Wolbachia* genes in an organism without maternal transmission. However, the *Wolbachia* sequences we identified were genetically close to those found in other Diptera, and we did not observe any signal of extensive divergence, which would be expected in the case of horizontal gene transfer^51,52^ (Figs 2 and 3).

Alternatively, mosquitoes could have been parasitized by *Wolbachia*-hosting nematodes^53,54^ or mites^55^. However, these *Wolbachia* strains belong to other, easily distinguishable supergroups (Fig. 2 and Fig. S1). Moreover, the analysis of *An. moucheti* F1 progeny confirms, at least in this species, that no other biological *Wolbachia* contamination was present in our analysis. In conclusion, our data suggest that *Wolbachia* is naturally present in the *Anopheles* species of Central Africa analysed in our study, and that is maternally inherited in *An. moucheti* (Table S2).

In Central African *Anopheles, Wolbachia* acquisition seems to be independent of the host phylogeny (Figs 2 and 3). Our results revealed that the genetic distances between *Wolbachia* and *Anopheles* are not positively correlated (Mantel test, p >0.05) (Fig. S2). Therefore, *Wolbachia* and the host lineage seem to evolve independently. The different larval ecology of these species suggests other ways of lateral transfer (e.g., during nectar feeding^39^). On the other hand, we found that all the *Anopheles* species belonging to the same complex shared related *Wolbachia* strains (Fig. 3). Permeable reproductive barriers among members of the same complex could facilitate the intermittent movement of the bacterium^56^. Interestingly, although they share similar *Wolbachia* strains, sibling species showed different infection prevalence. Indeed, *An. carnevalei* and *An. m. nigeriensis* exhibited frequencies lower than 15%, whereas *An. nili* and *An. moucheti*, their respective counterparts and the most important malaria vectors in their complex, displayed frequencies higher than 60% (Table 1). Moreover, our *An. gambiae* and *An. coluzzii* populations were infected by different *Wolbachia* strains than those detected in Burkina Faso and Mali. Similarly, in mosquitoes^36^ and ants^57^, the same species is infected by different *Wolbachia* strains according to the region. The availability of whole-genome sequences for *Wolbachia* strains^58^ will enlighten the intricate phylogenetic relationships among the different strains in *Anopheles*.

## Conclusions

*Wolbachia* has emerged as a biological tool for controlling vector-borne diseases^14,15^. In this study, we demonstrated the natural presence of this endosymbiont bacterium in a large number of *Anopheles* species, including the five major malaria vectors in Central Africa. It has been shown that *Wolbachia* ability to interfere with pathogen transmission depends on the bacterium strain. Therefore, our results offer the opportunity to determine whether the different Anopheles-infecting *Wolbachia* strains affect *Plasmodium* transmission and/or *Anopheles* reproduction. Indeed, the three most infected species *(An. moucheti, An. nili* and *An. vinckei*) play an important role in human (the first two) and non-human malaria transmission in the deep forest of Gabon^35,59^. Therefore, we could investigate *Wolbachia* positive^24^ and negative^60,61^ effects on the susceptibility to *Plasmodium* infection in their natural hosts. Moreover, the strongest effect on suppression of pathogen transmission or reproductive manipulation has been observed in *Wolbachia* transinfections^16,18,62–66^. Therefore, the availability of *Wolbachia* strains that infect natural *Anopheles* populations offers promising opportunities for experimental and theoretical studies in *Anopheles*, and also in other mosquito families that are vectors of other diseases, including *Ae. aegypti* and *Ae. albopictus.* In conclusion, our findings are merely the “tip of the iceberg” in the study of *Wolbachia* use to control vector-borne diseases, particularly in malaria.

## Methods

### Research and ethics statements

Mosquitoes were collected in Gabon under the research authorization AR0013/16/MESRS/CENAREST/CG/CST/CSAR and the national park entry authorization AE16008/PR/ANPN/SE/CS/AEPN. Mosquito sampling using the human-landing catch (HLC) method was performed under the protocol 0031/2014/SG/CNE approved by the National Research Ethics Committee of Gabon.

### Mosquito sampling and DNA extraction

Mosquitoes were collected in eight sites across Gabon, Central Africa, from 2012 to 2016 (Fig. 1, Table 1, Fig. S1). These sites included sylvatic (national parks) and anthropic habitats (villages and cities). Adult females were collected using CDC light traps, BG traps and HLC. Collected specimens were taxonomically identified according to standard morphological features^29,67^. Then, they were individually stored in 1.5 mL tubes at −20°C and sent to CIRMF for molecular analysis. When possible, at least 30 mosquitoes (from 1 to 58) for each *Anopheles* species from different sites were selected for genomic analysis. Total genomic DNA was extracted from the whole body using the DNeasy Blood and Tissue Kit (Qiagen), according to the manufacturer’s instructions. Genomic DNA was eluted in 100 μL of TE buffer. Specimens belonging to the *An. gambiae* complex, *An. funestus* group, *An. moucheti* complex and *An. nili* complex were molecularly identified using PCR-based diagnostic protocols^30–33,68^.

### *Wolbachia* screening and Multi Locus Sequence Typing (MLST) analysis

*Wolbachia* infection in adult females was detected by nested PCR amplification of a *Wolbachia-specific 16S* rDNA fragment (~400 bp) using 2 μL of host genomic DNA, according to the protocol developed in Catteruccia’s laboratory^24^ Amplification of this *16S* rDNA fragment in infected *Aedes albopictus* and *Culex pipiens* genomic DNA (data not shown) confirmed the performance of this nested PCR protocol to detect *Wolbachia* in many different mosquito species^24^. To detect potential contaminations, *Ae. albopictus* and *Cx quinquefasciatus* from Gabon were used as positive controls, and water and *Ae. Aegypti* as negative controls. Moreover, PCR amplifications for each species were carried out independently and in different days. The amplicon size was checked on 1.5% agarose gels, and amplified *16S* rDNA fragments were sent to Genewiz (UK) for sequencing (forward and reverse) to confirm the presence of *Wolbachia*-specific sequences. The DNA quality of all samples was confirmed by the successful amplification of a fragment (~800 bp) of the mitochondrial gene *COII* in all the *Anopheles* species under study^34,69^ PCR products were run on 1.5% agarose gels, and *COII* fragments from 176 mosquito specimens of the 25 species were sequenced (forward and reverse) by Genewiz (UK) for the *Anopheles* phylogenetic studies. *Wolbachia-positive* genomic DNA samples (2 μL/sample) were then genotyped by MLST using three loci, *coxA (~450 bp) ftsZ (~500 bp)* and *fbpA (~460 bp)*^70^, and according to standard conditions^44^ If the three fragments could not be amplified, a newly-developed nested PCR protocol was used. Specifically, after the first run with the standard primers, 2 μL of the obtained product was amplified again using internal primers specific for each gene: *coxA* (coxA_NF-2: 5‘-TTTAACATGCGCGCAAAAGG-3’; coxA_NR-2: 5‘-TAAGCCCAACAGTGAACATATG-3’), *ftsZ* (ftsZ_NF-2: 5‘-ATGGGCGGTGGTACTGGAAC-3’; ftsZ_NR-2: 5‘-AGCACTAATTGCCCTATCTTCT-3’), and *fbpA* (fbpA_NF-1: 5‘-AGCTTAACTTCTGATCAAGCA-3’; fbpA_NR-1: 5‘-TTCTTTTTCCTGCAAAGCAAG-3’). Cycling conditions for *coxA* and *ftsZ* were: 94°C for 5min, followed by 36 cycles at 94°C for 15s, 55°C for 15s and 72°C for 30s, and a final extension step at 72°C for 10min. For *fbpA*, they were: 94°C for 5min followed by 36 cycles at 94°C for 30s, 59°C for 45s and 72°C for 90s, and a final extension step at 72°C for 10 min. The resulting fragments (coxA, 357bp; *fbpA*, 358-bp; and *ftsZ*, 424-bp) were sequenced bidirectionally by Genewiz (UK). The new sequences obtained in this study were submitted to GenBank (accession numbers XX to XX, Supplementary Table 1). Unfortunately, the other three MLST genes (gatB, *wsp* and *hcpA)* could not be amplified, due to technical problems (i.e. multiple bands)

### Phylogenetic and statistical analysis

All *Wolbachia* sequences for the *16S, coxA, fbpA* and *ftsZ* gene fragments and for *Anopheles* COII were corrected using *Geneious* R10^71^. The resulting consensus sequences for each gene were aligned with sequences that represent the main known *Wolbachia* supergroups obtained from GenBank (see Table S1). Only unique haplotypes for each species were included in the analysis (haplotype was defined as a unique allelic profile for each examined locus). Inference of phylogenetic trees was performed using the maximum likelihood (ML) method and RAxML^72^ with a substitution model GTR + CAT^73^ and 1000 bootstrapping replicates. Finally, all MLST *Wolbachia* sequences were used to build phylogenetic trees using RAxML (GTR+CAT model, 1000 bootstrapping replicates). Trees were visualized with iTOL v.3.4.3^74^

To quantify the accuracy of the observed *Wolbachia* infection prevalence, the influence of sample size on its estimation was assessed. For this, it was assumed that *Wolbachia* prevalence within a host species followed a beta binomial distribution^27^ yielding many species with a low or a high *Wolbachia* prevalence but few with an intermediate one. This allowed quantifying, for each sample size, the proportion of samples (over 1,000 realizations) that could yield an estimate that was not significantly different from the prevalence over the whole population with a z-test and a significance threshold at 95%. As expected, sample size could be small for very low or very high prevalence (60 individuals are enough in 95% of cases for these extreme prevalence values), while it was much higher for intermediate prevalence values (up to 150 individuals for a prevalence value close to 50%).

All statistical analyses were performed using “R” v3.2.5 (R Development Core Team, http://cran.r-project.org/), with the addition of the “ggplot2” library^75^.

## Author contributions

D.A., O.D. and C.P. designed the experiments. D.A., O.A., N.R. and P.K., performed the experiments. D.A., F.S., C.C., O.D and C.P. analysed the data. D.A. performed the sequencing analysis. D.A. and B.R. performed the statistical analysis. D.A., N.R., M.N., F.M., B.M., C.P. and F.P. provided samples for the analysis. D.A., O.D. and C.P. wrote the manuscript.

## Supplementary Materials

**Text S1**. Mosquito taxonomic and molecular identification.

**Table S1**. Mosquitoes screened in this study and their accession numbers.

ID: Specimen identification name; Sites: Collection locations; Species: Morphological and molecular identification; COII_hap: COII haplotype for each species; COII: accession number for the cytochrome oxidase subunit II gene; 16S: accession number for the 16S rRNA gene; ftsZ: accession number for the filamenting temperature-sensitive mutant Z protein; fbpA: accession number the fructose-bisphosphate putative aldolase protein; coxA: accession number for the cytochrome c oxidase subunit I.

**Table S2**. Summary of the mosquitoes collected and screened in this study

**Table S3**. *Anopheles moucheti* F1 used to estimate vertical transmission.

**Figure S1**. Rooted maximum likelihood phylogeny of the filarial *Wolbachia* sequence isolated from one *An. coustani* specimen.

The tree was inferred with RAxML^72^ using the sequence of the filarial *COII* fragment amplified from the *An. coustani* specimen BNG78 (in blue) and public sequences (NCBI) and rooted with *Brugia malayi* as outgroup. The black dot on the branch indicates a bootstrap value >70% from 1000 replicates.

**Figure S2**. Scatterplot showing the genetic distances between *Wolbachia* strains *(16S)* and infected *Anopheles* species *(COII*).

Genetic distances were estimated as the number of different bases between the sequences of each pair of infected *Anopheles* specimens. The smoothed conditional mean (blue line) and the 95% confidence intervals (grey area) were plotted using the smoothing “gam” function of the ggplo2 library^75^.

**Figure S3**. Probability of detecting *Wolbachia* infection.

The probability was estimated for each sample size and infection prevalence value. The probability of correct estimation follows a black-blue gradient. The dashed line indicates the 90% probability of infection detection.

